# Mitigating the lack of knowledge about long noncoding RNA: extracting biological functions

**DOI:** 10.1101/258343

**Authors:** Yagoub A. I. Adam, Evandro Eduardo Seron Ruiz, Alessandra Alaniz Macedo

**Affiliations:** Inter-institutional Graduate Program on Bioinformatics, Department of Computer Science and Mathematics, FFCLRP, University of São Paulo, Ribeirão Preto, SP, Brazil; Department of Computer Science and Mathematics, FFCLRP, University of São Paulo, Ribeirão Preto, SP, Brazil.

**Keywords:** Long noncoding RNA, Biological Function, Information Extraction, Natural Language Processing

## Abstract

Long non-coding RNAs (lncRNAs) play crucial roles in diverse biological processes. For example, they help to regulate gene expression and to shape the 3D organization of the cellular nucleus. However, their functions in these processes are not well known. Most of the ongoing research efforts regarding lncRNAs have focused on predicting their properties and their functions. Here, we aimed to extract the biological functions of lncRNAs referred in the biomedical literature. To this end, we performed gene enrichment tests for 417 verified gene names extracted from the scientific literature. We also succeeded in extracting biologically significant functions from 2455 single sentences of 2513 scientific abstracts retrieved from the PubMed digital library. These sentences had to be filtered to eliminate those that did not mention a biological function. They were also split into 2-3-grams and the meaningful function annotations based on Gene Ontology terms were extracted. The results provided by the gene enrichment test and the knowledge gained by automatic extraction of information from the scientific literature showed that lncRNAs play a critical role in all biological and cellular processes. Although simple natural language processing techniques fulfilled our purposes of extracting concepts concerning the biological functions of lncRNAs, in the future, we aim to use more sophisticated methods, such as grammatical rules, to extract more functions and to generate a full database for the biological functions related to lncRNAs.

## 1 Introduction

According to the central dogma of gene expression, RNA molecules mediate the information transfer between DNA and proteins. Nevertheless, RNA molecules have been recently discovered to perform other biological functions: ribozymes can act as enzymes, and RNA can participate in the translation machinery [6]. Furthermore, RNA molecules with regulatory functions have been discovered [39], which is further evidence of the diverse functions of RNA molecules and the importance of RNA in complex biological functions. Several studies have revealed that a large amount of RNA is transcribed, but not translated into protein [39, 30, 35, 34, 29]. These types of RNA are defined as noncoding RNA; hence, they are transcribed from noncoding regions. The five major groups of noncoding RNA are:

1. Small nuclear RNAs (snRNAs), which catalyze RNA splicing;
2. Micro RNAs (miRNAs). They regulate the level of targeted gene expression and have characteristic length of between 22 and 26 nucleotides;
3. Small interfering RNAs (siRNAs), which inhibit the transcription of targeted genes with complementary sequences and have epigenetic regulation functions such as defense against infections and chromatin remodeling [3]. These RNAs typically display between 20 and 25 nucleotides and form double-stranded RNA molecules;
4. Piwi-interacting RNAs (piRNAs). They play a role in epigenetic regulations. The length of these RNAs ranges from 24 to 31 nucleotides [16];
5. Long non-coding RNA (lncRNAs). Numerous studies have revealed that many lncNAs containing over 200 nucleotides are transcribed. These diverse noncoding RNAs are classified as lncRNAs [40, 5]. LncRNAs take part in countless processes, including evolutionary functions, regulation of biological functions, stem cell differentiation, regulation of the cell cycle, and regulation of a variety of cellular pathways. Recent studies have shown that lncRNAs are emerging as important regulators of disease processes such as cancer [30, 29, 14, 13, 28, 4, 25, 38, 2, 27, 21‒24].

As lncRNAs have a critical role in all the biological and cellular processes, many efforts have been made to characterize and to annotate lncRNAs. lncRNAs have been considered as part of the “dark matter of the genome” due to the limited understanding of their functions and activities. Understanding the functions of lncRNAs has become a holy grail in the field of bioinformatics and molecular biology, and new approaches to annotate lncRNAs are necessary.

In this study, we aim to investigate the role lncRNAs play in all cellular functions by using natural language processing (NLP) methods to further understand their role in genome as its is presented in the biomedical literature. We also intend to mitigate the lack of knowledge about lncRNAs by extracting information from specialized biomedical literature.

The remaining sections of this paper are organized as follows: Section 2 presents some related work in textual processing of biomedical information. Section 3 describes the proposed method to mitigate the lack of knowledge about lncRNAs. This method extracts biological functions from the biomedical literature. Section 4 discusses the results obtained. Section 5 presents our final remarks and suggestions for future work.

## 2 Textual Processing and Extraction

There are versatile approaches to extract interesting biomedical information from the literature [7]. These approaches mostly include:

– Text mining to named entity recognition (NER) and terminology extraction;
– Text classification; and
– Relationship extraction and hypothesis generation.

As our aim is to correlate lncRNA and their biological functions in the literature, co-occurrence of targeted terms in sentences is a simple method to extract targeted sentences. This method provides wide coverage of the sentences; however, it also extracts high numbers of unrelated sentences. As a result, this approach provides high sensitivity and low specificity for the targeted sentences [11].

On the other hand, the extraction of information by targeted patterns (also called pattern-based techniques) increases the specificity but lowers the sensitivity [36]. Pattern-based techniques are either supervised techniques, in which domain experts manually define patterns through analysis of reliable literature, or unsupervised techniques, whereas patterns are automatically defined and learned through well-annotated literature. To increase the accuracy of the information extracted by the pattern-based approach, the grammatical structure of the sentences should be considered [1].

A well-known NLP method to extract information from biomedical literature is the shallow parsing of sentences into word groups, also called n-grams [15, 20]. The outputs from parsers may be processed by either a supervised rule-based learning method [26, 8, 17, 33] or by an unsupervised learning approach, such as a Support Vector Machine [12, 19, 18].

## 3 Material and Method

To extract biological functions of lncRNAs from the biomedical literature we designed a pipeline that encompasses the following steps: (i) Abstract text retrieval from PubMed; (ii) Extraction of official gene symbols for gene enrichment test; (iii) Processing of the retrieved primary raw data set and the extraction of relevant sentences; and (iv) Extraction of biological functions from relevant sentences. Figure (1) summarizes this proposal.

**Fig. 1.**
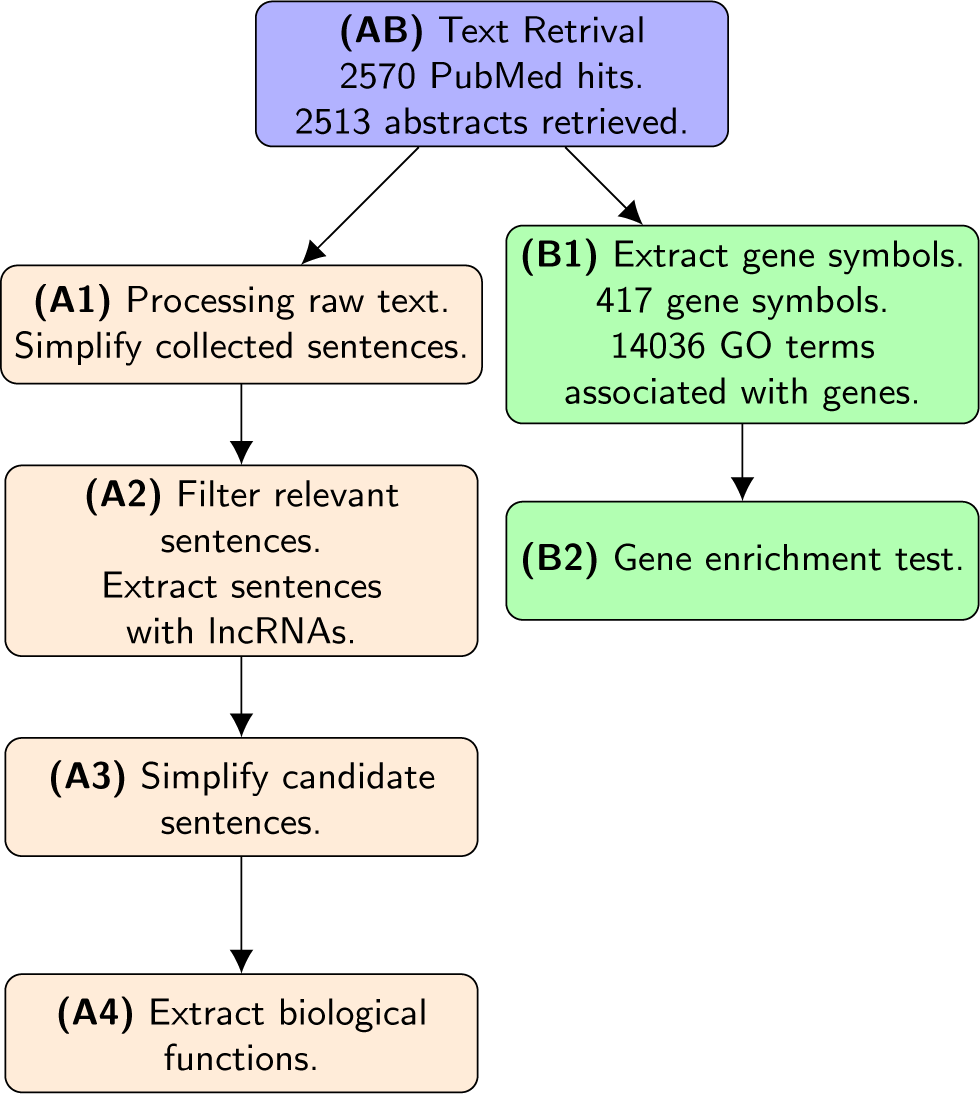
Flowchart showing the pipeline of our research method.

### 3.1 Text Retrieval from PubMed

The first pipeline task was the retrieval of relevant abstracts from PudMed by querying with the term ‘noncoding RNA [All Fields]’ as a single query term in order to achieve higher sensitivity and a high recall. See Figure 1, **AB**.

### 3.2 Extraction of Gene Symbols and Gene Enrichment Test

To extract the gene names from the texts, we noticed that sentences enclosed in parentheses and words between three and six characters were extracted as they show a potential of being gene symbols. Afterward the Gene Ontology (GO) terms related to the gene symbols were verified using the g:rProfile web-tool version r1536_e83_eg30 [32, 31]. The Gene Ontology enRIchment anaLysis and visuaLizAtion tool (GOrilla) was also used to identify and visualize the enriched GO terms for the verified genes [10, 9]. See Figure 1, **B1** and **B2**. Figures 2 depict the results that are commented in Section 4.

**Fig. 2.**
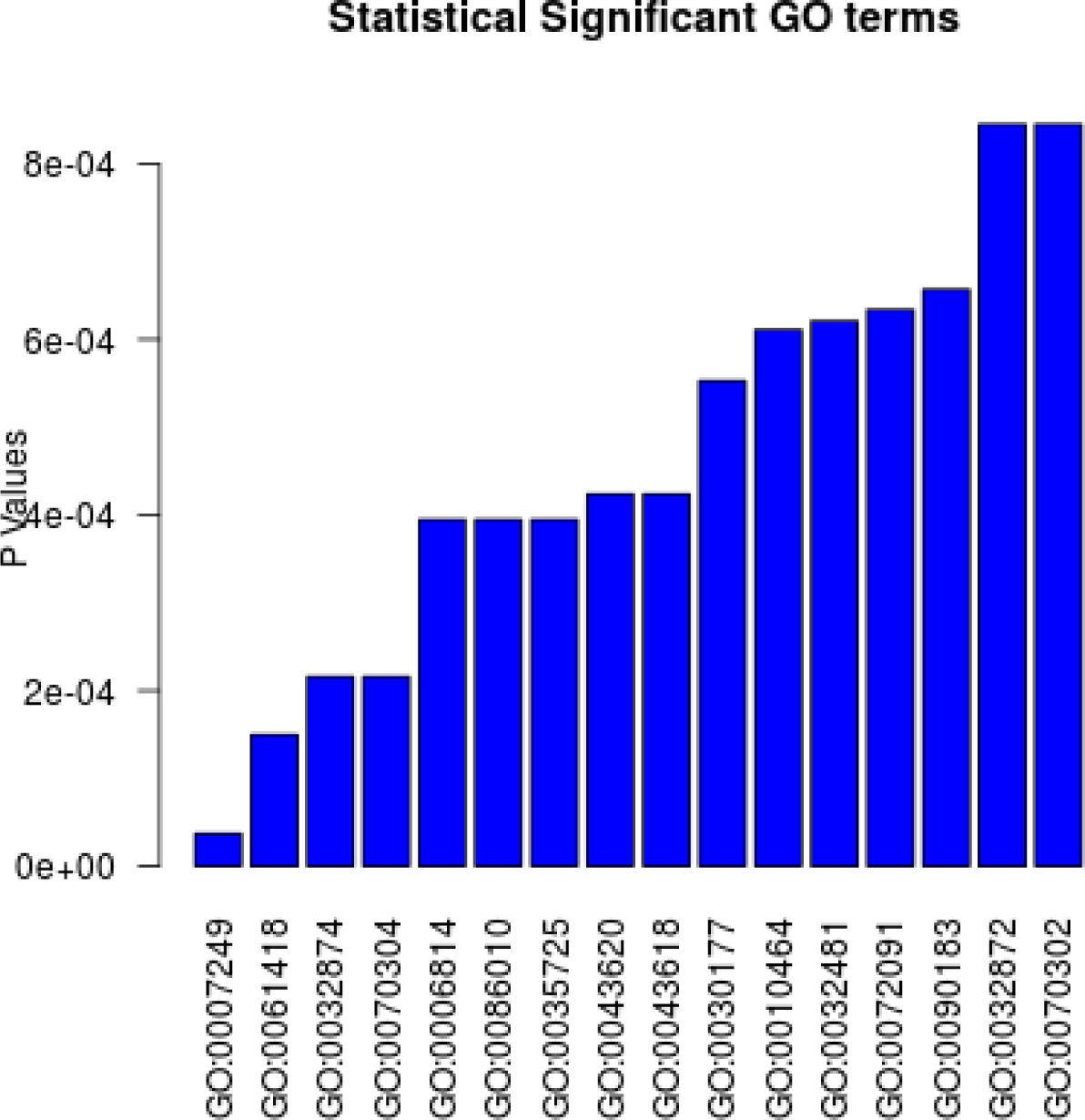
Significant GO terms from the gene enrichment test.

### 3.3 Processing of the Primary Retrieved Text Raw Data

At this stage, the main target was to simplify the collected sentences by eliminating the potential errors that could occur during the following steps. The text processing task for error prevention consists of:

– Conversion of all characters to lowercase;
– Removal of parenthetical remarks, which release the terms inside a pair of parentheses;
– Normalization of compound words and extended sentences containing the signs ‘+’,‘-’, or ‘” into single words by removing non-letter characters;
– Replacement of multiple spaces with single space; and finally
– Replacement of the synonymous terms that refer to lncRNAs with a new defined abbreviation. See Table (1).

**Table 1.**
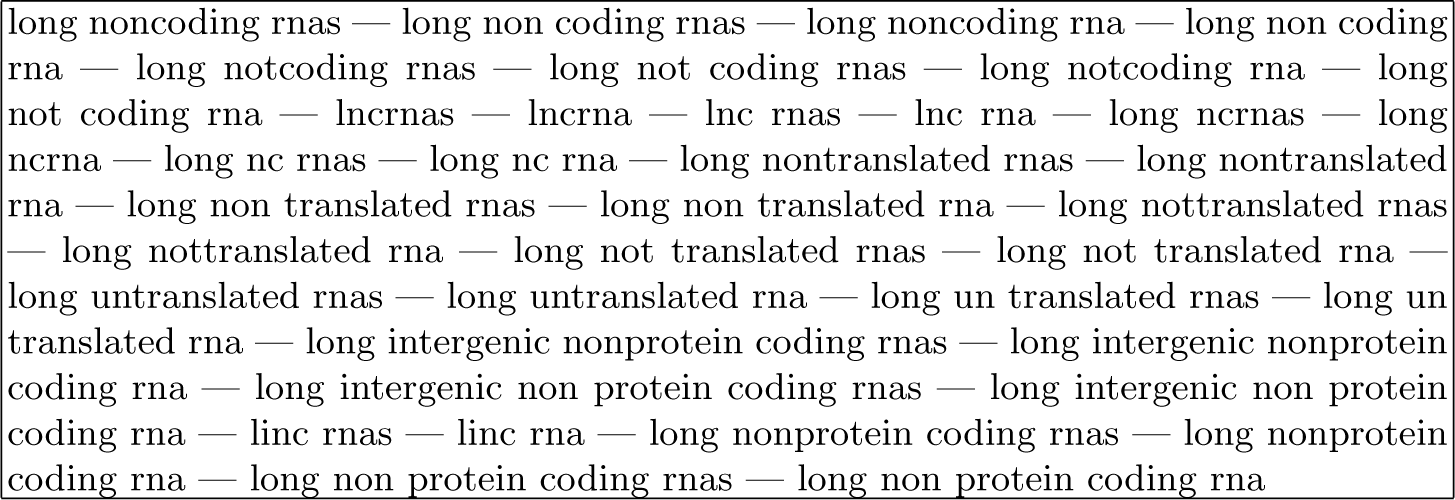
Predefined lncRNAs synonyms which replaced with single term ‘lncRNAs’.

**Table 2.** Description of the selected GO terms.

| GO Term | Description |
| --- | --- |
| GO:0007249 | I-kappaB kinase/NF-kappaB signaling |
| GO:0061418 | regulation of transcription from RNA polymerase II promoter in response to hypoxia |
| GO:0032874 | positive regulation of stress-activated MAPK cascade |
| GO:0070304 | positive regulation of stress-activated protein kinase signaling cascade |
| GO:0006814 | sodium ion transport |
| GO:0086010 | membrane depolarization during action potential |
| GO:0035725 | sodium ion transmembrane transport |
| GO:0043620 | regulation of DNA-templated transcription in response to stress |
| GO:0043618 | regulation of transcription from RNA polymerase II promoter in response to stress |
| GO:0030177 | positive regulation of Wnt signaling pathway |
| GO:0010464 | regulation of mesenchymal cell proliferation |
| GO:0032481 | positive regulation of type I interferon production |
| GO:0072091 | regulation of stem cell proliferation |
| GO:0090183 | regulation of kidney development |
| GO:0032872 | regulation of stress-activated MAPK cascade |
| GO:0070302 | regulation of stress-activated protein kinase signaling cascade |

See Figure 1, **A1**.

### 3.4 Extraction of relevant sentences

The abstracts were split into sentences and the relevant sentences were selected. The relevant sentences were defined as sentences that contained a predefined abbreviation of long noncoding RNA, according to Table 1 and which were extracted for further analysis. See Figure 1.

### 3.5 Extraction of the biological functions of lncRNAs from relevant sentences

From the relevant sentences, eventual lncRNAs biological functions were extracted from up to 10 words that came after the keyword ‘lncRNA’. This was accomplished by shallow parsing the collection of words into bi-grams and trigrams. See Figure 1, **A4**.

## 4 Results and Discussion

Querying the PubMed search engine using “noncoding RNA [All Fields]” as the query term, our method retrieved 2513 abstracts out of 2570 related PubMed Id (see Attachment S1 Query result). The remaining 57 abstracts failed to be retrieved mainly due to problems such as format issues: abstracts were in PDF or as image (or other non-XML format), or the documents had no abstract section, such as laboratory notes, clinical records, patents, etc.

The gene enrichment task was conducted for 417 verified genes by using the Gorilla tool. The results of this test showed that lncRNAs are associated with genes present in larger amount in various important biological processes such as:

– Oncogenesis or tumorigenesis;
– development and differentiation of stem cells as well as regulation of stem cell proliferation;
– Regulation of biological processes (such as transcription) that also regulate the immune system through positive regulation of the production of Type I Interferon;
– Communication within cellular environment through signaling pathways and transport of celular materials; and
– Response to external and internal stress.

To extract the biological functions of lncRNAs from the collected abstracts, the first step was to extract 2455 sentences as candidate sentences containing biological functions. Second, the proposed method collected up 10 words that followed the predefined lncRNA keywords as a final text. This final text was filtered by removing non-relevant high frequency terms. Bigrams and trigrams were then filtered, revealing that lncRNAs are associated with various biological processes such as:

1. Cancer initiation/inhibition, cancer development, cancer metastasis, and cancer progression for various types of cancers including: bladder cancer, gastric cancer, lung cancer, prostate cancer, breast cancer, colorectal cancer, pancreatic cancer, cervical cancer, thyroid cancer, esophageal cancer, liver cancer, colon cancer, urothelial cancer, and brain cancer, cancer growth, cancer invasion, adenocarcinoma, urothelial carcinoma, esophageal carcinomas, neuroblastoma, squamous carcinomas, pancreatic gliomas, lymphoma, lung carcinomas, hepatoblastoma, hepatocellular carcinomas, melanoma, adenocarcinoma, colorectal carcinomas, meningiomas, and glioma vascular cancer;
2. Regulation of biological processes such as: gene expression and transcription, chromatin remodeling and interactions, and regulation of genome;
3. Cellular processes like cell growth, cell migration, cell proliferation, and cell differentiation; and
4. Response to external and internal stress through cell signaling and response to diseases.

Figures 2, 3, and 4 and the table in the S2 attachments depict our results. Figure 2 shows the the *p*-values of 16 statistically significance GO terms. These terms are related to lncRNA functions. These significant GO terms were considered associated with biological function as they presented *p*-values ≤ 0.05, according to a hypergeometric distribution applied using the GOrilla software, as explained in subsection 3.2. See Table 4 for the list of the 16 GO terms. Refer to Table S2 in the attachments for more details about the gene enrichment task.

**Fig. 3.**
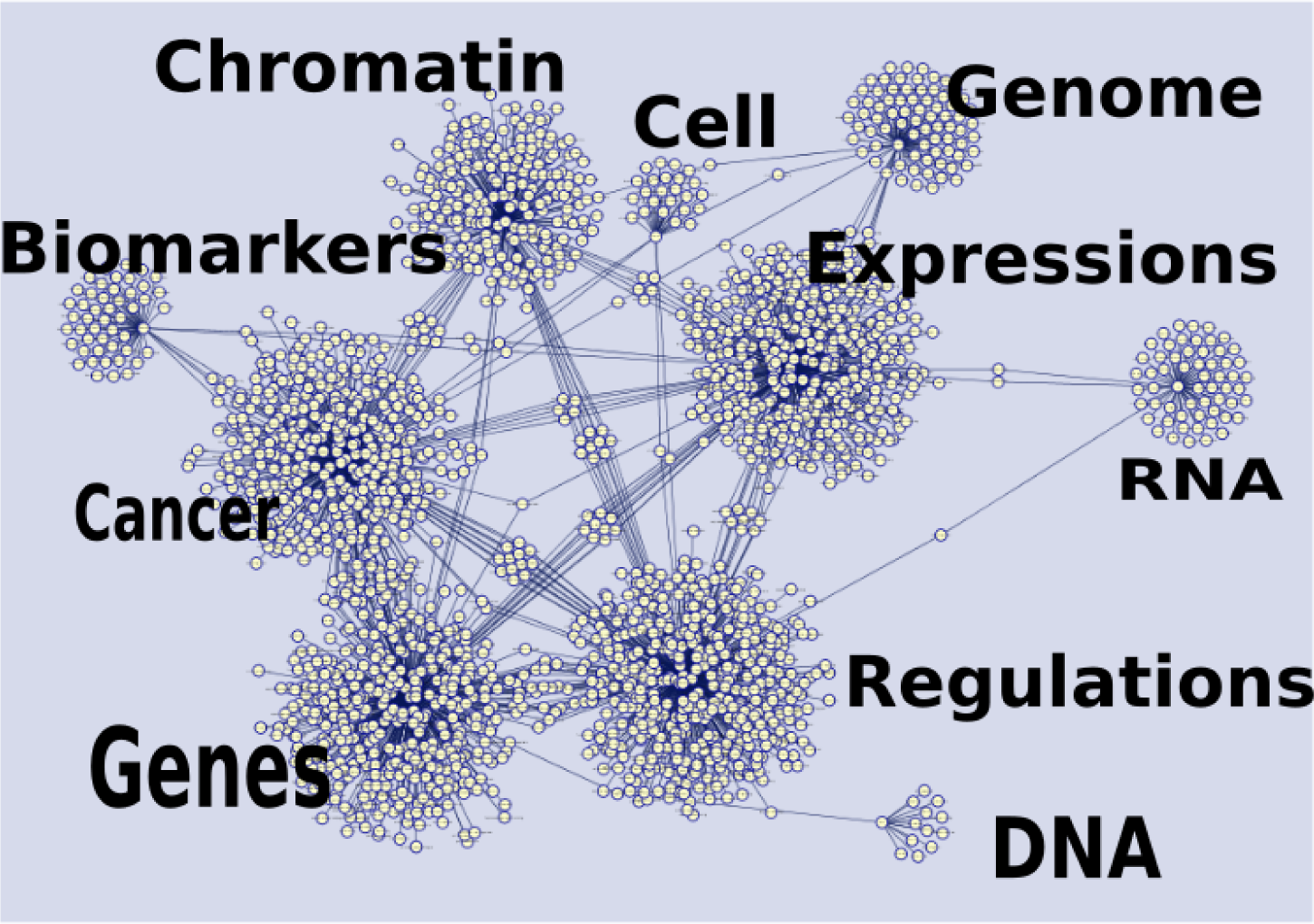
Networks of bigrams terms in the final text using DNA, Genes, Cancer, Biomarkers, RNA, Expressions, Regulations, Cell, and Chromatin as nodes.

**Fig. 4.**
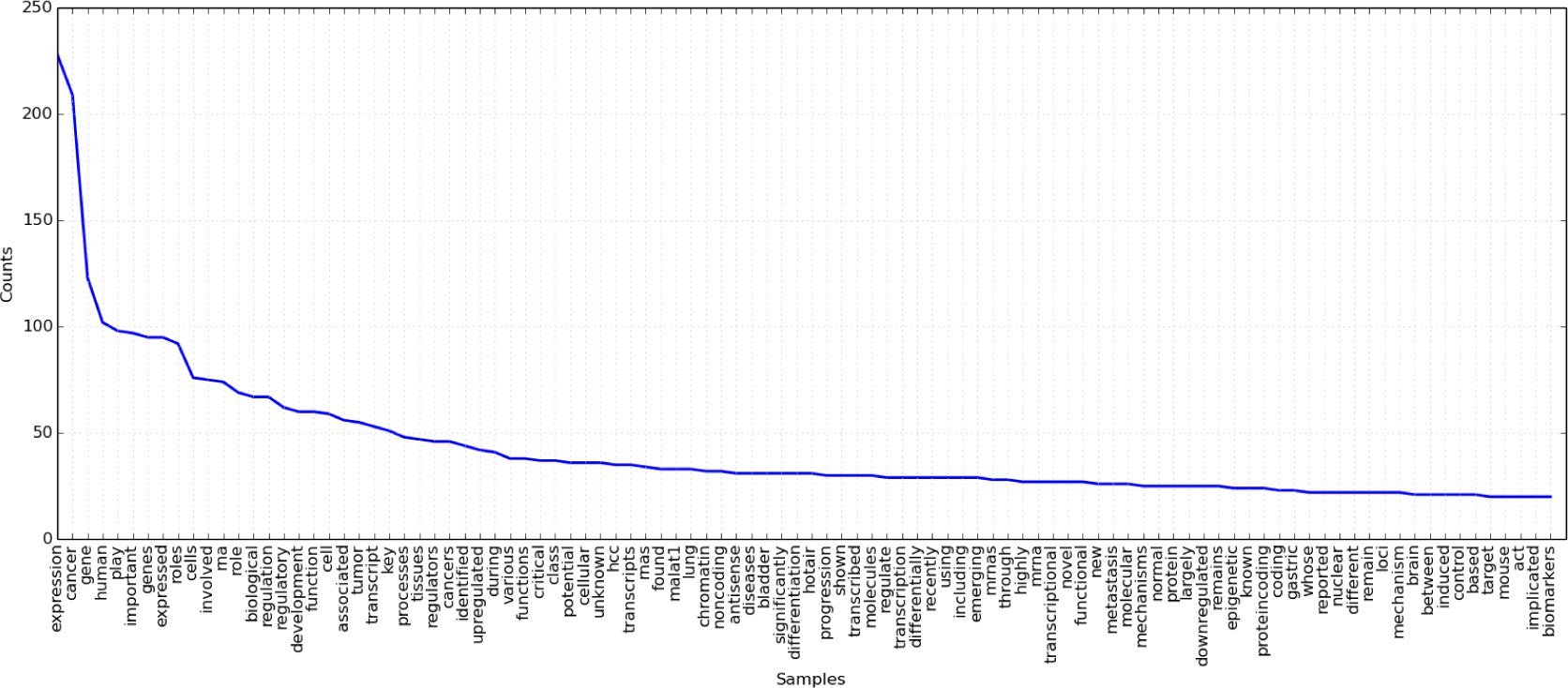
The 100 most frequency words in the final text.

Figure 3 shows the network of biologically significance bi-grams around one of the following terms: Chromatin, Cell, Genome, Expressions, RNA, Regulations, DNA, Genes, Cancer and Biomarkers. As one of these terms appears in a relevant sentence, obtained as explained in Section 3.4, all the co-occurring bi-grams were collected and linked sequentially as they were found. The bi-gram network formed by this procedure was visualized using Cytoscape [37]. This figure reinforces the number of possible significant functions associated with lncRNAs.

Now considering only single terms, words, Figure 4 shows the 100 top most frequent words in the relevant sentences, i.e. sentences which contain the lncRNA predefined term. Although many words in this list may be considered very ordinal words, by the looking closer for biologically meaningful words the general roles of lncRNAs within the cell could be predicted.

## 5 Final Remarks

The amount of scientific literature available on health subjects is always increasing. For instance, PubMed currently comprises over twenty million MEDLINE citations. Despite the positive effects of having a large amount of information available, some negative aspects arise. The huge volume of scientific information burdens professionals because searches for accurate information are complex and time consuming.

This paper is an initial effort to demonstrate how we have started to discover knowledge by automatically extracting information from a scientific digital repository and, more specifically, extracting terms related to lncRNAs biological function. We have used simple NLP methods to extract concepts related to the biological functions of lncRNAs and to extract genes associated with lncRNAs. We have found that RNA molecules not only function as mediators but also have more complex biological functions. lncRNAs are emerging as important regulators of cellular process and functions and disease processes. Further investigations with more elaborate natural language processing methods, such as the use grammatical rules, might help to extract even more functions and knowledge related to lncRNAs.

## 6 Acknowledgments

Yagoub A. I. Adam gratefully acknowledges the financial support of CAPES.

## A Attachments

All the following attachments can be found in goo.gl/oxK1HX

1. S1 Query result in XML format
2. S2 Table of significant GO terms
3. S3 Annotated Genes

